# Navigating the oligomeric landscape of the periplasmic stress response protease-chaperone DegP with charge detection mass spectrometry

**DOI:** 10.64898/2026.08.21.746289

**Authors:** Kent R. Vosper, Aizaz Humayun, Matthew S. McPartlan, Ashleigh Wint, Madison Turner, Siavash Vahidi, Evan R. Williams, Robert W. Harkness

## Abstract

DegP is a periplasmic protease-chaperone essential for protein quality control and virulence factor trafficking in Gram-negative bacteria. In its apo form, DegP adopts a dynamic ensemble of oligomers derived from trimer building blocks through two competing self-assembly pathways. Upon engaging client proteins, apo DegP oligomers redistribute into discrete cage structures inside which the clients are encapsulated. The cage ensembles formed depend on the size of the bound clients, and notably can include 12mers, 24mers, and 60mers. Previous studies mapped the DegP oligmeric landscape using dynamic light scattering, analytical ultracentrifugation, nuclear magnetic resonance spectroscopy, and electron cryomicroscopy, modalities which in general report ensemble averages and often cannot directly delineate closely related coexisting species. Here, we apply charge detection mass spectrometry (CDMS) to directly measure the masses of individual DegP ions, resolving the complete oligomeric distribution in the absence and presence of four clients of increasing size. We reveal previously undetected odd-numbered oligomers and quantify the relative abundance of each assembly. Through heat-cool cycling CDMS experiments, we track cage distribution changes and reveal client protection and refolding, providing a direct view of the chaperone capabilities of DegP. These results further establish CDMS as a powerful single-molecule tool for dissecting heterogeneous protein assembly landscapes.

## Introduction

Protein homeostasis in the bacterial periplasm is maintained by a network of chaperones and proteases that ensure the proper folding, trafficking, and recycling of client proteins.^1,2^ Among these, DegP is a widely conserved dual protease and chaperone from the High temperature requirement A (HtrA) family that is essential for bacterial survival under stress conditions including heat,^3,4^ oxidative,^5^ and osmotic shock^6^ (Figure 1A). DegP is additionally implicated in virulence factor trafficking in the ESKAPE pathogens, where it escorts virulence factors through the periplasm to the outer membrane in a partially unfolded state for secretion (Figure 1A).^7,8^ The involvement of DegP in bacterial protein homeostasis and virulence has made it a promising target for antibiotic development,^9,10^ where modulatory ligands that either inhibit or enhance its activity could disrupt bacterial function and prevent infections.

**Figure 1.**
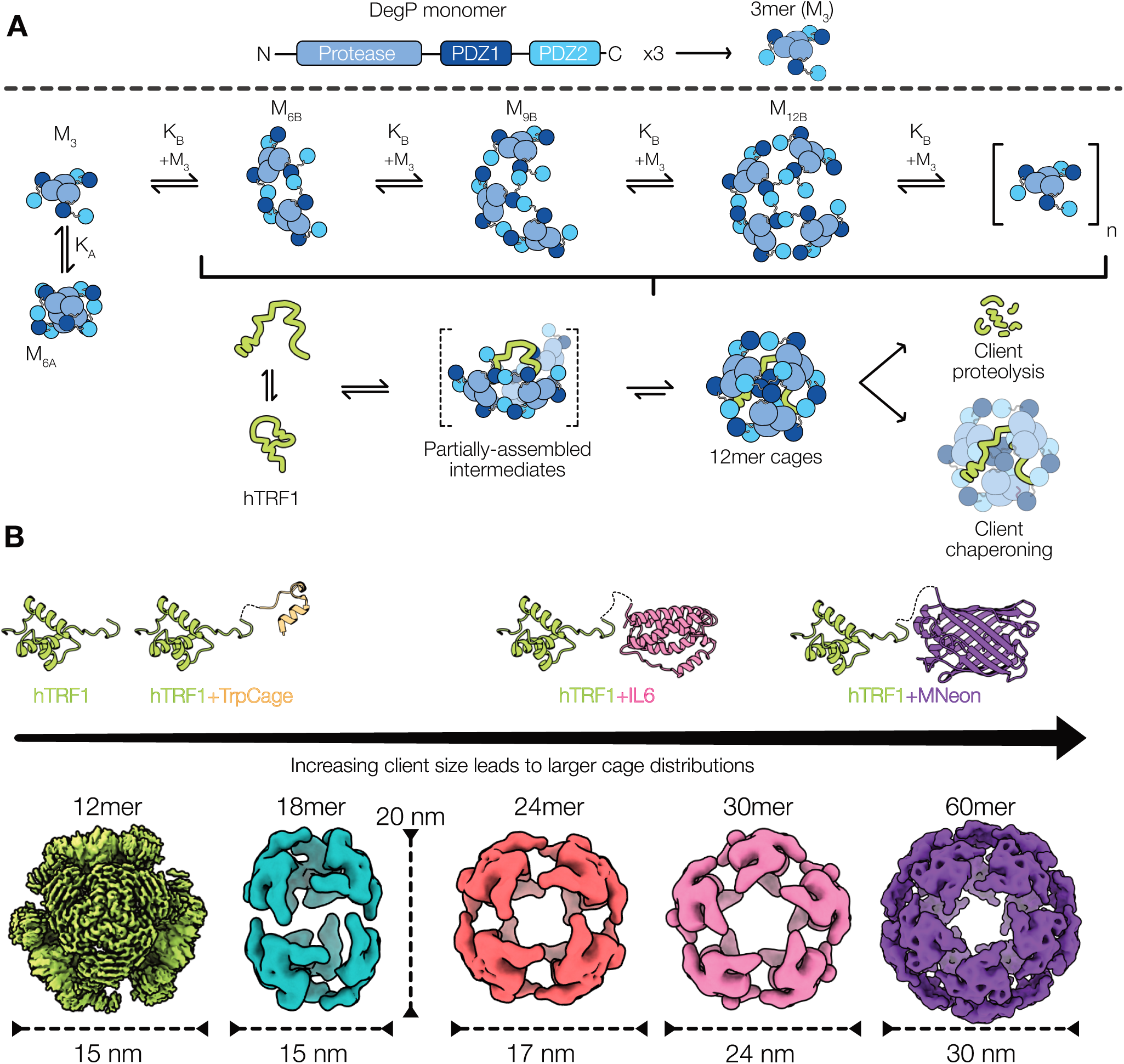
The DegP assembly landscape. (A) DegP monomer architecture (top): each monomer consists of a protease domain (light blue) followed by two PDZ domains, PDZ1 (dark blue) and PDZ2 (cyan). Three monomers associate into the trimer building block (M_3_) that gives rise to the higher-order assemblies. In the apo state, trimers assemble into larger species through two competing pathways, A and B (middle). In path A, trimers adopt the canonical hexamer state, M_6A_, observed in crystal structures,^13,30^ according to K_A_. In path B, trimers oligomerize through an isodesmic mechanism governed by K_B_ into a broad distribution of higher order species (M_6B_, M_9B_, M_12B_, …, M_3nB_). In the presence of hTRF1, the apo ensemble is reorganized into bound 12mer cages through partially assembled intermediate states (bottom). The 12mer cages can either proteolyze or chaperone the client. (B) Structural models of the DegP clients used in this study (top) with the corresponding cryo-EM density maps of DegP:client cages obtained in earlier studies (bottom). Cages are colored to match clients that produce them and are arranged left to right by client size to illustrate that increasingly large cage distributions are formed in response to them. The smallest model client is the DNA binding domain of hTRF1 (PDB 1ITY), which binds tightly to DegP. This has been fused to the C terminus of TrpCage (PDB 1L2Y), IL6 (PDB 2IL6), and MNeon (PDB 5LTR) as an affinity tag to generate the chimeric clients of increasing size that lead to progressively larger cage assemblies. DegP bound to hTRF1 (green) or TrpCage-hTRF1 (orange) predominantly forms 12mer cages (EMD-28781), IL6-hTRF1 (pink) mixtures of 18mer (EMD-28800), 24mer (EMD-ZZZ), and 30mer cages (EMD-28806), and MNeon-hTRF1 (purple) 60mer (EMD-28808) and larger cages.

The functional unit of DegP is a trimer built from monomers consisting of a serine protease domain followed by two tandem PDZ domains (hereafter referred to as PDZ1 and PDZ2, Figure 1A top). We recently showed that apo DegP trimers further assemble into a broadly populated distribution of higher order oligomers, governed by two competing pathways (path A and path B, Figure 1A middle).^11^ The formation of this ensemble is sensitive to DegP concentration, solution temperature, and ionic strength.^12^ In path A, DegP adopts a “resting” hexamer, formed primarily by protease-protease’ domain interactions between two stacked trimers (prime indicates domains from the partner trimer). This sequesters the trimer active sites, thereby preventing spurious proteolysis of substrates.^13^ In path B, DegP adopts large and labile cage-like species mediated by edge-wise intertrimer PDZ1:PDZ2’ domain interactions, following an isodesmic growth mechanism.^13–15^ Under physiological temperature (37 °C) and protein concentration (∼20 and ∼200 μM in resting and stress conditions respectively),^11,12^ the oligomeric landscape of DegP encompasses species from both pathways, including trimers, path A hexamers, and a broad distribution of large path B assemblies (Figure 1A middle).^11,12,16^ Upon client engagement, this pre-organized cage-like ensemble converts into a new distribution consisting of discrete cages where clients are encapsulated in the cage interiors (Figure 1A bottom, Figure 1B). These cages, like the apo species, form through the intertrimer PDZ domain contacts which are bolstered by contributions from the bound substrate.^11,12^ These cages can range from 600 kDa tetrahedral 12mers to 3 MDa icosahedral 60mers, and in some cases can be as large as subcellular organelles (Figure 1B).^11^

We have established the DegP self-assembly landscape through a combination of dynamic light scattering (DLS), analytical ultracentrifugation (AUC), nuclear magnetic resonance (NMR) spectroscopy, and electron cryomicroscopy (cryo-EM).^11,12,16^ Although these ensemble biophysical techniques have provided detailed insights into DegP oligomer formation and client binding, each possesses inherent limitations related to sample preparation, the solution conditions that can be explored to map self-assembly landscapes, and difficulties in resolving individual states when applied to polydisperse oligomeric systems.^17–19^ Critically, none of these methods can simultaneously and directly measure the masses and populations of all coexisting oligomeric species in a single experiment. Charge detection mass spectrometry (CDMS) offers a fundamentally different approach to characterizing heterogeneous macromolecular assemblies.^20,21^ In CDMS, individual ions generated by native electrospray ionization are confined in an electrostatic trap and passed repeatedly through a charge-detection tube. Each pass of an ion through the tube induces a signal with an amplitude proportional to the charge on the ion (z). The ion frequency of oscillation, combined with dynamic ion energy measurements,^22^ provides the mass-to-charge ratio (*m/z*). Particle mass is thus obtained directly from the simultaneous and independent measurements of *m/z* and z for each individual ion trapped.^23–25^ Because the mass of each ion is individually measured, CDMS produces mass histograms without ensemble ion averaging.^20^ This makes CDMS particularly well suited for studies of large, heterogeneous mixtures in the 100 kilodalton (kDa) to gigadalton (GDa) range, especially where the overlapping charge-state envelopes of a broad and heterogeneous distribution can no longer be resolved by conventional native mass spectrometry.^21^ The technique has been successfully applied to study viral capsid assembly intermediates,^26^ proteasomal complexes,^27,28^ and synthetic nanoparticles,^29^ thereby establishing it as a powerful tool for studying polydisperse macromolecular systems.

Here, we apply CDMS to map the oligomeric distribution of DegP in its apo form and in the presence of four client proteins of increasing hydrodynamic radius (Figure 1B). We reveal previously undetected odd-numbered oligomers (e.g. 15mer and 21mer) that represent labile apo assemblies and partly constructed client-bound cages. Through quantitative analyses of our CDMS data, we extract oligomer population fractions for the DegP landscape, including states that are populated to only a few percent. Using heat-cool cycling experiments we study DegP assembly in simulated bacterial heat shock conditions where DegP is overexpressed to capture and process substrates. These temperature cycling investigations reveal redistribution of cage species, protection of clients from aggregation, and refolded client egress, providing a direct view of DegP’s chaperone function. Our study orthogonally validates the two-pathway DegP assembly model and demonstrates the unique advantages of individual-ion mass measurement for studying dynamic protein assemblies.^11,12,16^ The capabilities that we demonstrate here further highlight CDMS as a valuable tool for rapidly dissecting heterogeneous oligomeric landscapes.

## Results

### CDMS resolves the apo DegP oligomeric distribution

To characterize the oligomeric landscape of apo DegP by CDMS, we used DegP purified under denaturing conditions and subsequently refolded to remove background client proteins that can induce undesired cage formation.^12,30^ The DegP construct implemented here harbors the catalytically inactivating S210A substitution to prevent autoproteolysis and allow for studies of client binding in the absence of degradation. It also includes C57S/C69S substitution that removes Cys residues which could form spurious inter-trimer crosslinks and complicate the interpretation of CDMS data. In what follows we refer to this construct as DegP.

We electrosprayed apo DegP at 20 µM monomer concentration in 200 mM acetate buffer at pH 7 and 25 °C and measured *m/z* and z values of individual ions (Figure 2A). These solution conditions were selected to ensure that, prior to electrospray, DegP adopted a broad mixture of path A and path B oligomers for testing the capabilities of CDMS in analyzing heterogeneous mixtures of DegP assemblies.^11,12,16^ The ion count as a function of *m/z* and z, and the associated mass histogram, revealed a series of well-resolved peaks (Figure 2B, C). We ascribed these peaks to specific DegP oligomers based on their masses, in multiples of trimers (Figure 2C). We note that the width of each peak in these datasets likely includes contributions from oligomeric states that are not strictly multiples of trimers (i.e. containing monomers and dimers), as well as the adduction of salts, solvent, or small molecules and not the intrinsic resolving power inherent to the CDMS instrument.^22–24,31^ Hereafter we analyze the mass histograms in terms of trimer units, as these are the most stable forms of DegP assemblies.^12,16^

**Figure 2.**
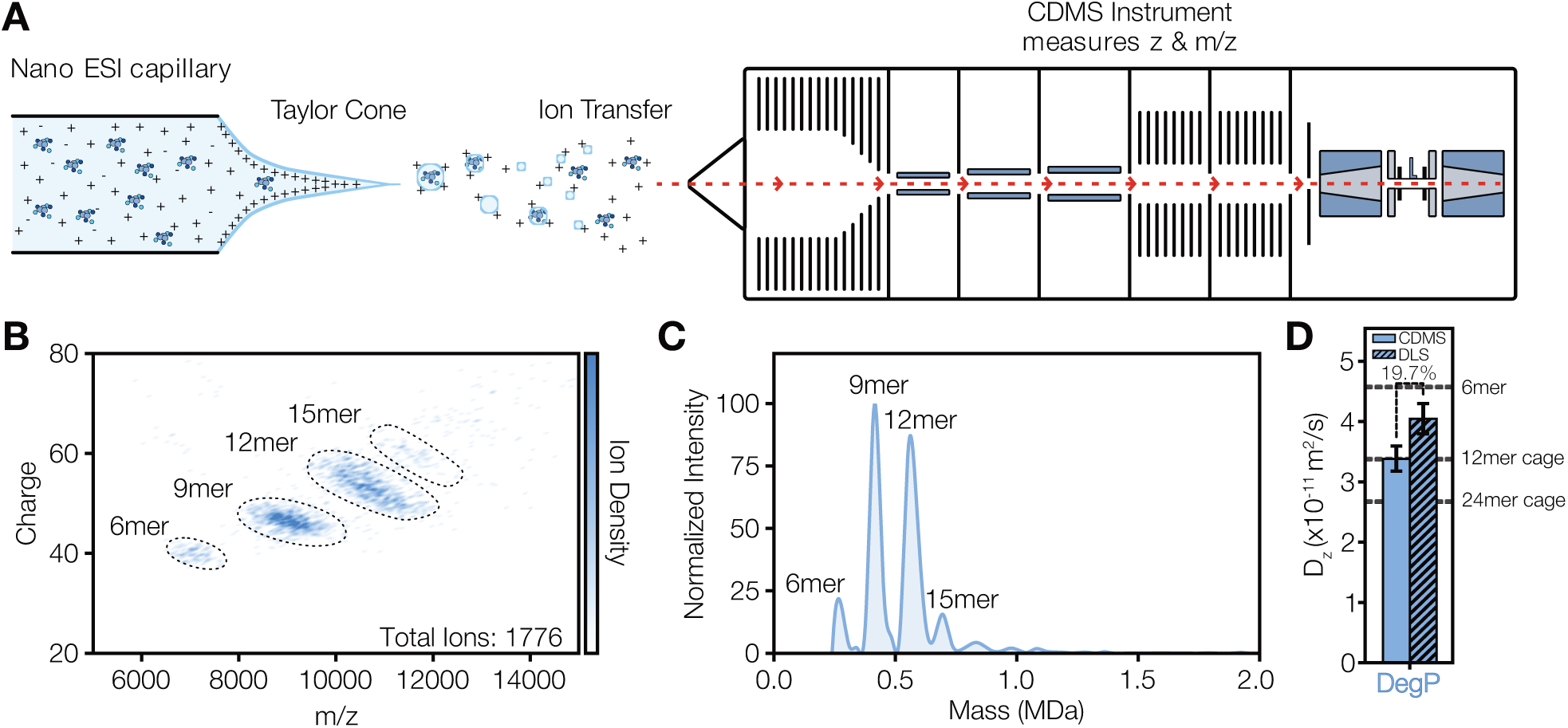
CDMS of apo DegP. (A) Schematic representation of the CDMS instrument used in this study. Protein molecules are electrosprayed and both *m/z* and z data for individual ions are measured, from which mass histograms are generated. (B) z vs. *m/z* of apo DegP (25 °C). Data points corresponding to individual ions are colored from white to blue, indicating ion density. Ion distributions for 6, 9, 12, and 15mers of apo DegP are encircled with black dashed ovals. (C) Kernel density estimates of mass histograms of apo DegP oligomers generated from the *m/z* vs. z plots in (B), normalized by the maximum intensity. (D) D_z_ calculated from the mass histogram in (C) (solid blue bar). The DLS-derived value (hatched blue bar) has been reproduced from Harkness et al., 2021 for comparison.^12^

We quantified the relative population of each species using a deconvolution procedure (see SI Methods, Figure S1, Table S1), in which a series of Gaussians describing the relative contributions of discrete states to the mass histogram are fit to the dataset. The dominant peaks of the mass histogram were assigned to the apo hexamer (M_6_, ∼280 kDa 7.5%), 9mer (∼420 kDa, 38%), 12mer (M_12_, ∼560 kDa, 42%), and 15mer (M_15_, ∼700 kDa, 7%). Small amounts of 18mer (M_18_, ∼840 kDa, 3%) and 21mer (M_21_, ∼980 kDa, 1%) were also present. Of the 1,776 ions that were weighed, 51 ions had masses above 1.0 MDa, indicating that less than ∼3% of the sample was in the form of even higher-order states. No appreciable trimer was identified in the CDMS data, highlighting the fact that the ensemble is shifted to higher-order states under the solution conditions we employed.

The presence of these larger species is consistent with prior temperature- and concentration-dependent DLS and AUC measurements.^11,12,16^ However, in those cases, the estimates of fractional populations of assemblies within the apo oligomer distribution were extracted through fitting of mass action models to multidimensional datasets, while here we could quantify them directly. To more robustly compare our results here with the z-average diffusion constants (D_z_) extracted from prior DLS measurements,^11,12^ which are measures of the effective oligomer ensemble size,^12,19,32,33^ we leveraged the population fractions of the apo DegP species measured using CDMS to estimate D_z_ values (see SI Methods, Figure 2D). We note that the DLS-derived D_z_ values have been reproduced here from prior investigations for clarity.^11,12^ As expected, the D_z_ value calculated from the CDMS data for S210A/C57S/C69S DegP was somewhat smaller than that from DLS (∼20%) for S210A DegP measured under similar experimental conditions (25 °C and 20 µM DegP monomer concentration; sodium acetate rather than sodium phosphate was implemented here for optimal signal-to-noise) (Figure 2D). This reflects the additional C57S/C69S substitution we used to simplify our CDMS analyses. Substitutions in this region of DegP, the protease domain LA loop, destabilize the path A hexamer,^12,16^ allowing for a greater abundance of the larger path B oligomers and thus a reduction in D_z_. Taken together, our CDMS measurements of the apo DegP ensemble confirm the two-pathway assembly model in which trimers associate into hexamers via path A and into higher-order oligomers via path B.^12^

### CDMS maps redistribution of DegP into client-bound cage ensembles

Having established that CDMS faithfully reports on the apo DegP oligomeric ensemble, we next used it to examine distributions of client-bound cage structures that are formed through reshaping of the apo DegP energy landscape upon client engagement (Figure 1A, B).^11,16^ To this end, we employed a series of model substrates of increasing hydrodynamic radius that we used in prior DLS and cryo-EM studies of client-bound DegP cages.^11,16^ The size of the cage ensembles formed in the presence of these clients is positively correlated with client size and can extend to multiple MDa assemblies, providing a wide dynamic range for CDMS studies of DegP. The smallest substrate consists of the DNA binding domain of the human telomere repeat binding factor 1 (hTRF1, r_h_ ∼1.5 nm),^11,34^ which binds to the protease and PDZ1 domains of DegP with high affinity (stoichiometric binding).^11,16^ The larger substrates are chimeric constructs containing N- and C-terminal “domains”. The N-terminal portions of the chimeras are formed by the TrpCage miniprotein, IL6, and MNeon respectively.^35–37^ The C-terminal segment consists of hTRF1, which serves as an affinity tag for binding to DegP. In the cages formed by these clients, the N-terminal portion is projected into the interior of the cages, tethered by the hTRF1 DegP anchor. The size of the N-terminal domain of the chimera controls expansion of the cages through extension and reorganization of DegP’s flexible PDZ domains and interdomain linkers.^11^ In what follows, we refer to these chimeric constructs as TrpCage-hTRF1 (r_h_ of 1.9 nm), IL6-hTRF1 (3.0 nm), and MNeon-hTRF1 (3.7 nm).^11^

DegP (20 μM subunit concentration) and each of the four clients (10 μM) were mixed, equilibrated, and electrosprayed for CDMS measurements. We selected the 2:1 ratio of DegP:client to approximate stress conditions where DegP is overexpressed and therefore can be present in excess relative to its substrates.^12,16^ This leads to partly filled cages that hold clients rather than degrading them.^16^ The CDMS data reveal a progressive shift in the oligomeric distribution toward larger cage species with increasing client size (Figure 3A, B), as anticipated.^16^ We quantified the relative abundance of each species in mass histograms by Gaussian peak fitting approach described above (SI Methods, Figure S1, Table S1). Given the widths of the measured peaks and spectral overlap, particularly for the larger cage ensembles obtained with IL6-hTRF1 and MNeon-hTRF1, we were only able to estimate approximate masses for the dominant effective particle sizes. We point out that, with the 2:1 DegP:client ratio used, the additional bound client masses within the cages are small relative to DegP (∼47 kDa monomer), particularly for hTRF1 (∼7 kDa) and TrpCage-hTRF1 (∼9 kDa). We could not discriminate between similarly sized cage architectures with different bound client stoichiometries, which can in principle be close in mass. Therefore, we used the known cage structures from earlier work as reference states for assigning client-bound peaks in our CDMS data.^11,30^

**Figure 3.**
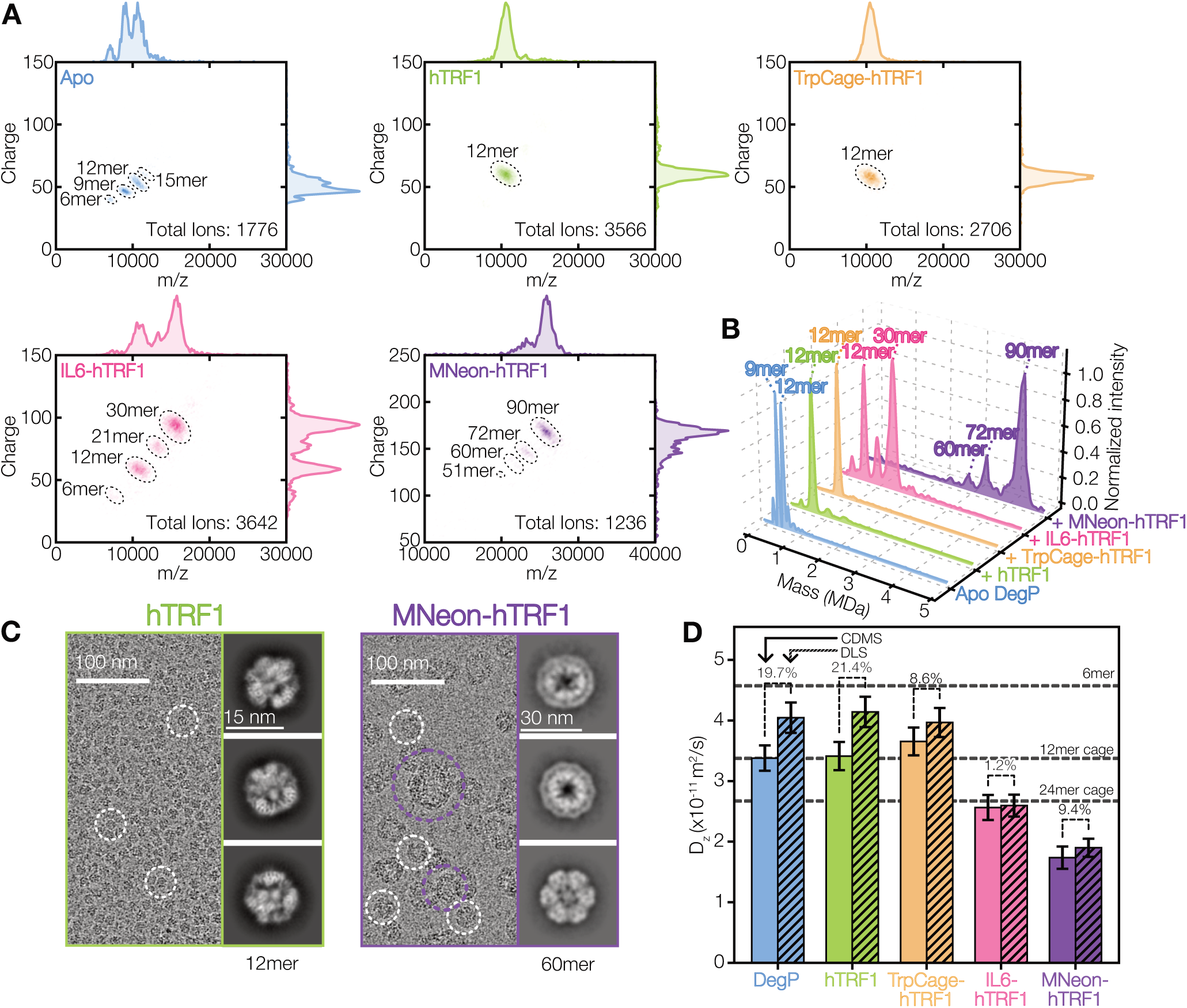
CDMS of DegP cage distributions as a function of increasing client size. (A) *m/z* to charge 2D kernel density estimates of mass histograms of apo and client-bound DegP structures. 1D projections of *m/z* and z spectra are displayed to the top and right respectively. Apo DegP (blue) assembles into labile pre-organized cage-like oligomers. In the presence of hTRF1 (green) and TrpCage-hTRF1 (orange), DegP primarily forms tetrahedral 12mers. IL6-hTRF1 (pink) leads to a distribution of DegP cages ranging from 6mer to 30mer, and MNeon-hTRF1 (purple) displays a distribution of larger cage assemblies from 51mers to 90mers. The dominant oligomeric species are encircled with black dashed ovals and labeled. (B) Kernel density estimates of mass histograms generated from CDMS of DegP in the presence of the clients. (C) Cryo-EM micrographs and 2D particle class averages of DegP cage formations in the presence of hTRF1 and MNeon-hTRF1. hTRF1-bound DegP primarily adopts 12mer structures, whereas MNeon-hTRF1-bound DegP forms a broad ensemble of cage states, including 60mers (circled in white), as well as amorphous larger species (circled in purple). Cryo-EM micrographs and 2D class averages have been reproduced from our prior study, Harkness et al., 2023.^11^ (D) D_z_ values calculated from CDMS distributions (solid colored bars) for apo DegP and bound to clients. DLS-derived values (hatched colored bars) measured in our earlier study^11^ are reproduced here for comparison. The percent differences between the CDMS- and DLS-derived D_z_ values are shown above each pair of bars (19.7%, 21.4%, 8.6%, 1.2%, and 9.4% for apo DegP, hTRF1, TrpCage-hTRF1, IL6-hTRF1, and MNeon-hTRF1, respectively). Horizontal dashed lines are the diffusion constants at 30 °C for the apo path A hexamer, a 12mer cage, and a 24mer cage, shown as visual references (see SI Methods). Errors (shown as 2 standard deviations) were obtained through Monte Carlo approaches.^66,67^

For the smallest clients, hTRF1 and TrpCage-hTRF1, (Figure 3A, B – green and orange traces respectively) the distribution is dominated by a 12mer cage peak (∼560 kDa, consistent with a tetrahedral cage, ∼86% for hTRF1, ∼100% for TrpCage-hTRF1; Table S1), with minor populations of 15mer (6%) and 18mers (4%). For IL6-hTRF1 (Figure 3A, B red), the dominant oligomers in the distribution are consistent with the tetrahedral 12mer and pentagonal bipyramidal 30mer cages (∼1.5 MDa). Strikingly, the mass histograms also contained substantial signal for previously unobserved 21mer cages, which may represent intermediate assemblies in the canonical octahedral 24mer and 30mer cage pathways. We note that the signature of the trigonal bipyramidal 18mer structure identified through cryo-EM is likely overlapped with that of the 21mer peak and is presumably present in significant amounts. A small fraction of a 6mer state (4.6%) was also resolved, possibly reflecting residual apo DegP or partly bound 6mer:IL6-hTRF1 species in equilibrium with the larger species observed. Finally, the CDMS data for DegP in the presence of the largest client, MNeon-hTRF1 (Figure 3A, B – purple) revealed mass peaks corresponding to extremely large assemblies. The dominant oligomers in the mass histogram are consistent with 72 and 90mers (∼14.5 and 77.3 % and ∼3.4 MDa and ∼4.2 MDa, respectively; Table S1, Figure S1), with trace amounts of 60mers (6.6%, ∼3 MDa) that we assigned to the icosahedral structure elucidated in earlier cryo-EM studies.^11^

We also compared the CDMS datasets collected in the presence of hTRF1 and MNeon-hTRF1, which serve as our boundary cases for the smallest and largest cage ensembles studied here, with qualitative particle distributions evaluated from cryo-EM micrographs (Figure 3C).^11^ The amounts of the 12mers in the hTRF1-bound sample that we measured in this study are in excellent agreement with those from our earlier investigations (Figure 3C, left). The relative amounts of the DegP:MNeon-hTRF1 species visualized by CDMS are also consistent with cryo-EM data collected on DegP in the presence of this client (Figure 3C, right), where 60mer and larger amorphous assemblies were found to coexist.^11^ We additionally calculated D_z_ values from the mass histograms obtained in the presence of each client for quantitative correlation with prior DLS-derived values^11,12^ (Figure 3D). For all clients studied here, the D_z_ values generated from the CDMS mass histograms were in good agreement with those extracted from DLS autocorrelation functions, with three of the four clients agreeing to within 10%.^11,12^ The largest difference was observed for hTRF1 (∼21% smaller CDMS *D_z_*), highlighting the contributions from the minor populations of the 15mer and 18mer in our CDMS conditions.^11^ CDMS measurements, therefore, robustly report on heterogeneous and broadly distributed DegP cage ensembles.

### Heat-cool cycling reveals reversible cage assembly, client protection, and client release

We next wondered whether CDMS could report on DegP’s chaperone function that assists in maintenance of protein homeostasis in the periplasm, and is implicated in the bacterial heat shock response.^3,4^ We therefore engaged in heat–cool cycling experiments monitored by CDMS. Prior to these measurements, we carried out thermal melting of DegP and each of the clients using circular dichroism to establish an appropriate experimental temperature at which the clients would be largely denatured while DegP remained folded (Figure S2). Each of the clients began unfolding at approximately 35 °C, while DegP did not show appreciable denaturation until roughly 60 °C. Only TrpCage-hTRF1 and hTRF1 were able to mostly refold during the cooling scan, while DegP, IL6-hTRF1, and MNeon-hTRF1 remained irreversibly denatured, suggestive of aggregation at high temperature. We therefore selected 50 °C for high temperature CDMS measurements, where DegP is safely folded while hTRF1 and TrpCage-hTRF1 are approximately 60% unfolded, and IL6-hTRF1 and MNeon-hTRF1 are roughly 40% unfolded (vertical dashed black line and colored circles in Figure S2). Importantly, the distinct responses of the small and large clients to heating and cooling enabled interrogation of how DegP cages shape their folding landscapes, either through protection from aggregation or assisted refolding, as read out via changes to CDMS distributions after temperature cycling.

Samples of 20 µM DegP and 10 µM client protein (2:1 ratio as above) were first prepared at ambient electrospray capillary temperature (30 °C for hTRF1 and TrpCage-hTRF1 and 35 °C for IL6-hTRF1 and MNeon-hTRF1 respectively), heated to 50 °C and then returned to 25 °C. At each stage, the samples were equilibrated for 10 minutes prior to acquisition of *m/z* and z. As anticipated, the initial distribution at 30 °C for hTRF1-bound DegP is dominated by the 12mer cage (Figure 4A, top left). Yet, strikingly, upon heating to 50 °C and cooling to 25 °C, the cage population redistributed, with the 12mer cages apparently disassembling into 6mers, leading to a roughly 1:1 ratio of 6mer:12mer, and a small population of trimers (Table S1, Figure S1). This thermal hysteresis suggests that some of the hTRF1 bound in the cages dissociated upon cooling via cage-induced refolding, causing partial collapse of the cage distribution to either apo or partly ligated 6mers and bound 12mers. The reversible circular dichroism-based thermal melt and annealing of hTRF1 strongly supports this conclusion (Figure S2). We note that this outcome is not likely to be due to destabilization of the cages with concomitant hTRF1 egress at 50 °C, as we have established in earlier NMR studies that hTRF1 is bound tightly within 12mers at this temperature via its unfolded state.^16^ We observed a similar phenomenon for TrpCage-hTRF1 (Figure 4A, top right). Notably, the new distributions for these clients obtained after temperature cycling did not match that for apo DegP we measured initially (Figure 2B, C), in keeping with the conclusion that roughly half of the hTRF1 and TrpCage-hTRF1 had emerged from the 12mer cages and was refolded.

**Figure 4.**
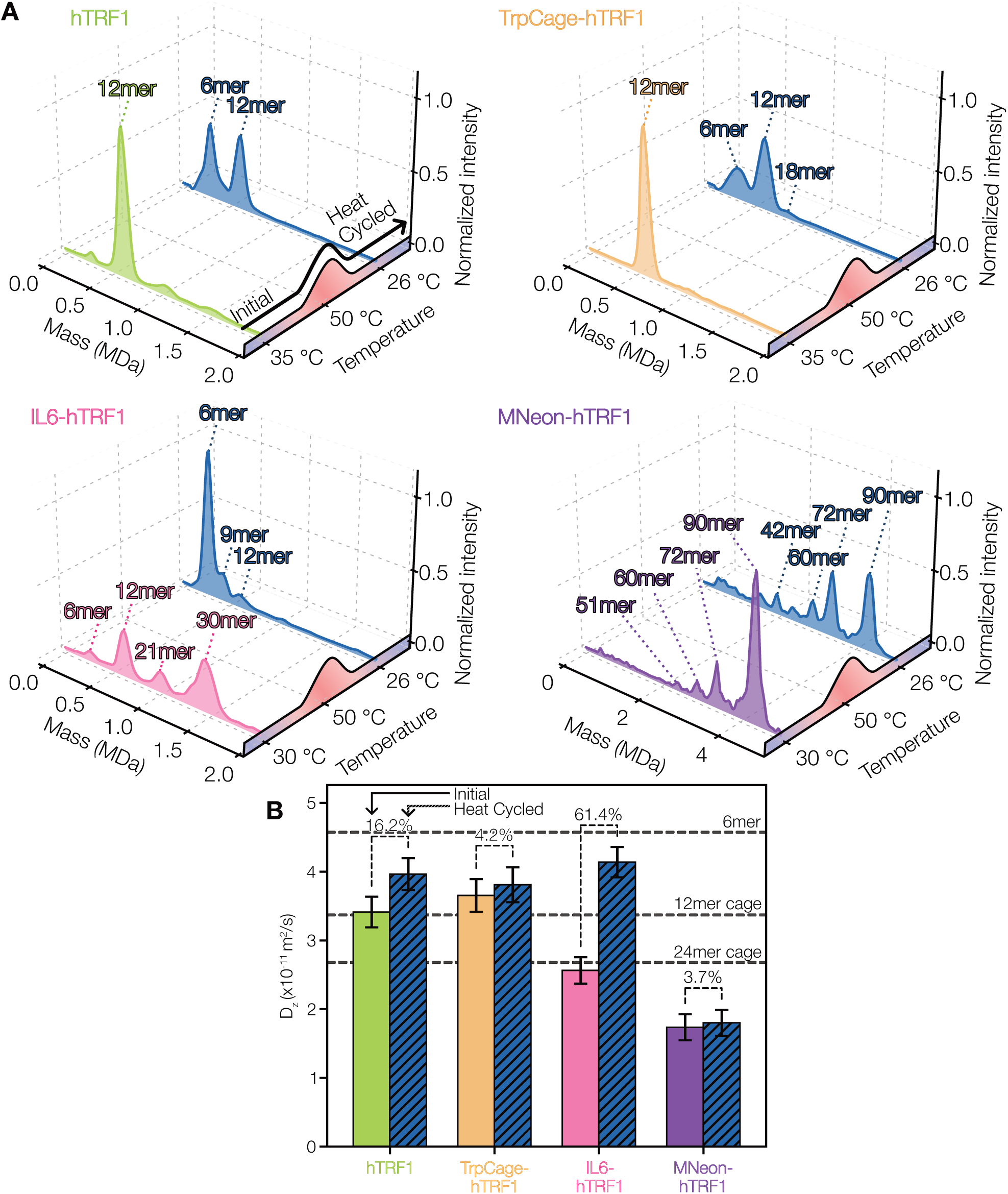
Monitoring DegP cage redistributions through temperature-dependent CDMS. (A) Kernel density estimates of mass histograms through heat-cool cycles for DegP in the presence of the four clients, normalized to the maximum intensity in each plot. Oligomeric species have been labelled both before and after heating. (B) CDMS-derived D_z_ values for hTRF1, TrpCage-hTRF1, IL6-hTRF1, and MNeon-hTRF1 measured at the initial temperature (client-colored solid bars) and after a heat–cool cycle (dark blue diagonally hatched bars). The percent change in D_z_ between the initial and heat-cycled conditions is shown above each bar pair (16.2%, 4.2%, 61.4%, and 3.7% for hTRF1, TrpCage-hTRF1, IL6-hTRF1, and MNeon-hTRF1, respectively). Horizontal dashed lines are the diffusion constants at 30 °C for the apo path A hexamer, a 12mer cage, and a 24mer cage, shown as visual references (see SI Methods). Error bars represent 2 standard deviations, estimated using Monte Carlo approaches.^66,67^

Heating and cooling of cages bound with IL6-hTRF1 led to the most dramatic outcome (Figure 4A, bottom left, Table S1, Figure S1). The initial distribution mostly featured 12mer, 21mer, and 30mer, shifting to nearly all 6mer accompanied by lesser populated 9mer and 12mer (15 and 8%, Figure 4A, bottom left). Considering the thermal melt data for this client, the temperature-cycled CDMS points to considerable aggregation despite being in the presence of DegP. This may in fact reflect temperature-induced weakening of the cages for IL6-hTRF1, causing high temperature egress and in turn unfolding and aggregation of the free client at 50 °C that cannot be rescued by DegP (Figure S2, see IL6-hTRF1 curve). One explanation for this effect could be that the IL6 “domain” of the chimeric client at high temperature has unfavorable interactions with other copies within cages, or with the PDZ1:PDZ2’ domain interfaces that form the cage architecture. Given that the apo DegP distribution (Figure 2B, C) was also not fully recovered in this case after cooling, as it would be if the entirety of IL6-hTRF1 was lost (free DegP remains folded at 50 °C), the mass histograms are, however, consistent with marginal protection from aggregation. Unexpectedly, in contrast to IL6-hTRF1, temperature cycling of the MNeon-hTRF1:DegP sample did not lead to a dramatic shift towards the apo DegP distribution (Figure 4A, bottom right). Instead, we observed substantial reorganization of the extremely large cage assemblies. Specifically, we measured an approximately 40% reduction in the initially observed 90mers, with concomitant increases in 72mers and 60mers (changing from 21 to 41% of the ensemble respectively), as well as the appearance of some smaller species including 42mers (6%). This indicates that most of the MNeon-hTRF1 persisted within cages at high temperature, as otherwise it would have fully irreversibly unfolded considering its apo thermal melt data (Figure S2, see MNeon-hTRF1 curves). Thus, MNeon-hTRF1 is strongly protected from aggregation by encapsulation within DegP cages.

To further quantify the extent to which the initial low temperature cage distributions were recovered, we calculated D_z_ values from the CDMS mass histograms obtained before and after temperature cycling of the DegP:client samples (Figure 4B). As expected, all clients except for IL6-hTRF1 had moderate increases in their *D_z_*values (∼16.2, ∼4.2, and ∼3.7% for hTRF1, TrpCage-hTRF1, and MNeon-hTRF1 respectively) toward the value for apo DegP (Figure 4B – see horizontal dashed line for the diffusion constant of the path A hexamer), reflecting the substantial fractions of cages remaining in these samples. IL6-hTRF1 had a much larger increase in D_z_ of approximately 61%, highlighting the dramatic shift of the ensemble to a nearly apo DegP distribution that we attributed to the extensive aggregation of this client.

## Discussion

The DegP protease-chaperone is critical to the growth and virulence of Gram-negative pathogens.^7,8,38^ It has emerged as a promising antibiotic target for mitigating or preventing infections, as demonstrated in relatively recent studies showing that its hyperactivation by tripodal peptidic ligands causes bacterial cell death.^9^ Understanding DegP’s self-assembly landscape, and how it responds to modulatory ligands, is therefore paramount to developing DegP-targeting compounds to combat infections, particularly in the face of ever-mounting antimicrobial resistance rates.^9,39^ Here we have validated CDMS as a key tool for investigating DegP oligomerization in apo and client-bound states, and through its heat shock cycle, covering the full spectrum of circumstances in which it may operate.

Our study reinforces the unique capacity of CDMS to quantify population fractions for distinct oligomeric states.^21,40,41^ For example, DLS studies report *D_z_* values for DegP + MNeon-hTRF1 that fall below the expected value for 60mer particles, suggesting either a well-defined larger state, or an average weighted towards species larger than 60mers.^11^ CDMS directly resolves the underlying distribution as a mixture of predominantly 60mer, 72mer, 90mer species, with minor populations of smaller as well as larger states (Figure 3A, B). Our temperature cycling results emphasize this limitation further (Figure 4A, B). Taking the MNeon-hTRF1 CDMS dataset as the most remarkable example (Figure 4A, bottom right), the marked reduction in 90mers and increases in 72mers and 60mers after temperature cycling, produced only a ∼4% increase in *D_z_* from its initial value. Evidently, a rearrangement of this extent is effectively invisible to DLS alone. CDMS profiles are therefore a vital complement to ensemble-average techniques including AUC, DLS, and NMR, which often struggle to decompose oligomeric ensembles into their constituent states, or are incapable of doing so.^11,12,16^ Although cryo-EM classification can in many cases discriminate between oligomeric states through its ability to classify particles that produce high resolution 3D reconstructions,^42^ this process is heavily biased and discards the vast majority of assemblies present. For this reason, it is not considered an analytical tool for quantifying particle populations.^28^ A key advantage of CDMS is the detection and quantification of intermediate and partially assembled species that may be lost during grid preparation, particle picking, or averaged into neighboring particle classes, in addition to the better defined states that cryo-EM captures.^11,26^ Consistent with this, CDMS detected significant populations of odd-numbered species in the presence of larger clients (Figure 3A), representing assemblies containing non-symmetric arrangements of trimers, which were not identified in prior cryo-EM studies of DegP.^11^

Self-assembling proteins regulate numerous biological processes across all domains of life, including cell division,^43^ motility,^44^ signalling pathways that control gene expression,^45^ and the formation of biomolecular condensates that modulate enzymatic reactions and protein folding,^46–48^ among many other essential functions.^49^ Perturbations to the oligomerization propensities of such proteins underlie neurodegenerative^50,51^ and neurodevelopmental disorders,^52^ cancers,^53,54^ and numerous other pathologies. Notable examples are the amyloid-forming proteins tau^50^ and Aβ,^51^ where it is now recognized that small early-stage oligomers drive cytotoxicity, rather than mature fibrils. CDMS has resolved these “hidden” species which are not visible with microscopy or ensemble methods.^55^ Moreover, it has allowed for an examination of how these neurodegenerative seeds form and are influenced by post-translational modifications.^55^ CDMS has also provided critical insights into the formation of capsid protein “nuclei” that seed formation of viral particles, and aggregration pathways of antibodies.^26,40,41,56^ Mechanistic studies of these of these types of systems invariably center on illuminating which oligomeric states are populated and how these ensembles respond to stimuli such as ligands or stress.^9,11,12,57^ The sensitivity of CDMS to shifts in oligomeric distributions detected as averages by other techniques opens the door to more deeply understanding how self-assembly is dysregulated in disease.

Our temperature-dependent studies of DegP assemblies showing cage reorganization and client aggregation emphasize the strengths of CDMS in analyzing protein conformational landscapes and thermal stability.^31,41,58^ Indeed, CDMS is finding applications in the development of thermostable biologics, particularly in the engineering and formulation of antibodies, which are typically concentrated to several hundred mg/mL for administering to patients and must survive variable transport conditions.^59–61^ These concentrations strongly favor the formation of high-molecular weight aggregates which can provoke severe immune reactions,^62^ and therefore these oligomers must be suppressed through rational design of antibody monomers with elevated stability. AUC is increasingly used in antibody development to identify and quantify discrete antibody oligomers but requires lengthy experiment times,^63,64^ while CDMS provides this information in a few minutes and can be used over a wide temperature range to assay traces of aggregation.^41^ Temperature-dependent CDMS measurements, as we have demonstrated here, therefore offer a direct and rapid strategy for assessing both protein folding and the stability of biologics in treatment development pipelines.^40,57^

The temperature cycling CDMS experiments illustrate the multi-well energy landscape governing DegP assembly established in prior studies.^11,12,16^ Notably the shifts in cage distributions upon cooling emphasizes the lability of the DegP landscape in reaction to client burden, where it shifts towards its apo distribution as the level of available substrates decreases. The partial release of clients suggests a mechanism by which DegP can sequester them during heat stress and subsequently release them upon return to lower temperatures, consistent with its chaperone function.^12,65^ This provides an opportunity for clients to refold rather than be degraded, thereby preserving the protein content of the periplasm (Figure 5). This work adds dimensionality to the picture of DegP that has emerged, one in which the plasticity of its trimers built from flexible domains and extensible interdomain linkers enables the rapid formation of dynamic oligomeric ensembles to respond to bacterial stress.

**Figure 5.**
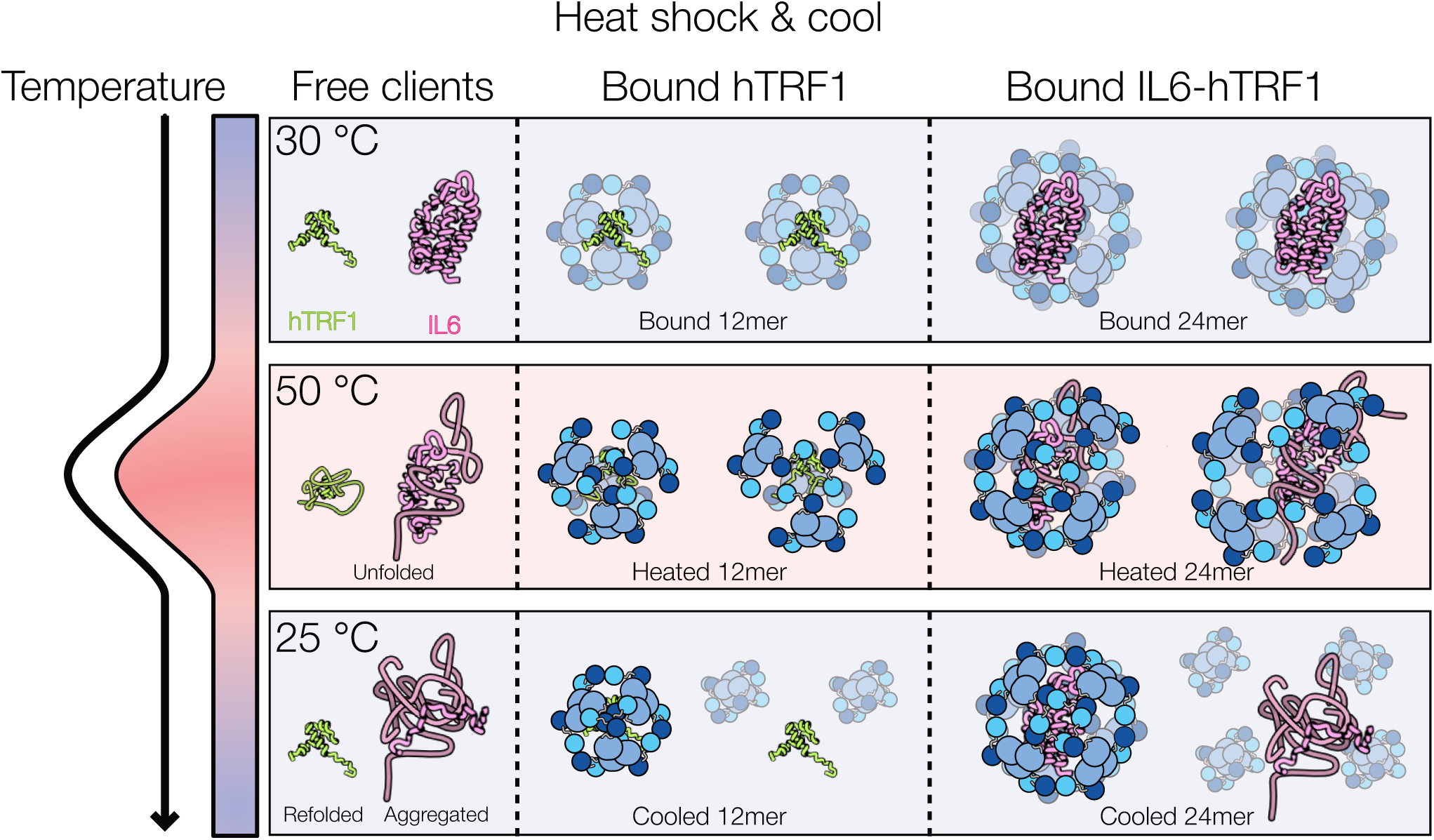
Model of DegP chaperoning clients through a heat shock cycle as established by CDMS. The free clients hTRF1 and IL6-hTRF1 (left column) and their complexes with DegP (middle and right column respectively) are shown at 30 °C (top row), 50 °C (middle row), and 25 °C (bottom row). The temperature gradient is indicated on the left. We have omitted the hTRF1 tag structure from IL6-hTRF1 in this schematic for simplicity. At 30 °C both clients in the absence of DegP are folded (top left). In the presence of DegP, DegP encapsulates hTRF1 within a 12mer cage (top middle) and IL6-hTRF1 within a 24mer cage (top right, taken as the average over the 12, 21, and 30mer IL6-hTRF1-DegP cages found here for simplicity). On heating to 50 °C the free clients largely unfold, with IL6-hTRF1 forming aggregates (middle left). When bound within the 12mer and 24mer cages, the clients are protected from aggregation despite increased thermal fluctuations of the cages at high temperature (expanded cages shown in the 50 °C row, middle and right). Note that one IL6-hTRF1:DegP cage is shown less tightly associated than the hTRF1:DegP complexes, with free IL6-hTRF1 emerging to depict the onset of its high temperature aggregation. On cooling to 25 °C free hTRF1 refolds, whereas IL6-hTRF1 remains irreversibly aggregated (bottom left). A fraction of the hTRF1-bound DegP cages release folded hTRF1 and form apo oligomers (bottom middle). While the majority of the IL6-hTRF1 aggregated at high temperatures, despite being in the presence of DegP, some IL6-hTRF1:DegP cages remain indicating partial protection from aggregation (bottom right).

## Acknowledgements

This contribution is dedicated to the memory of Prof. Wei Zhang. We are immensely grateful for his invaluable donation of research infrastructure that has facilitated these and other fruitful investigations. K.R.V acknowledges support from a Graduate Tuition Scholarship from the University of Guelph and an NSERC Doctoral scholarship. A.H. acknowledges support from an International Doctoral Tuition Scholarship from the University of Guelph. Financial support was provided by a Discovery Grant from the Natural Sciences and Engineering Research Council (NSERC) of Canada (RGPIN-2021-02843) to S.V., (RGPIN-2025-03947) to R.W.H., and (The National Science Foundation Division of Chemistry, grant number CHE-2506157) to E.R.W. The authors thank Dr. Casey J. Chen for assistance in acquiring mass spectrometry data.

## Notes

### Competing Interest Statement

The authors have declared no competing interest.

