## Supplementary Information for "Navigating the oligomeric landscape of the periplasmic stress response protease-chaperone DegP with charge detection mass spectrometry"

**This file contains:**

**Materials and methods**

**Table S1:** Populations of apo and client-bound DegP oligomers obtained from numerical fitting of CDMS kernel density estimate data.

**Figure S1:** Numerical fitting of CDMS kernel density estimates.

**Figure S2:** Thermal unfolding of apo DegP and clients monitored by CD spectroscopy.

**Materials and methods**

*Protein purification*

DegP S210A/C57S/C69S (lacking the catalytic serine to prevent auto and client degradation, and Cys resides to prevent spurious disulfide crosslinks; hereafter referred to as DegP), hTRF1, and the chimeric model clients TrpCage-hTRF1, IL6-hTRF1, and MNeon-hTRF1 were expressed and purified as described previously,^1^ with minor modifications. Each construct included an N-terminal His_6_-SUMO tag for affinity purification. Briefly, these proteins were expressed in T7 express *lysY* competent *Escherichia coli* (New England Biolabs). Cells were grown in LB broth incubated at 37 °C until an OD of ~0.6-0.8 was achieved. Protein expression was induced with 0.2 mM isopropyl-β-D-1-thiogalactopyranoside (IPTG). Protein expression was allowed to proceed for ~18 hours at 18 °C. Subsequently, cells were harvested via centrifugation, resuspended in lysis buffer (50 mM Tris-HCl, 6 M Gdn-HCl, pH 8.0 for DegP; 100 mM NaH_2_PO_4_, 500 mM NaCl, pH 8.0 for clients), and lysed by sonication. In general, each protein was purified according to nickel affinity chromatography,^2^ affinity tag cleavage and removal,^1^ an intermediate polishing chromatography step in the case of DegP and MNeon-hTRF1 (hydrophobic interaction chromatography using a SkillPak TOYOPEARL Butyl-650M column (Tosoh) for DegP, or ion exchange with a HiTrap SP Fast Flow 5 mL column (Cytiva) for MNeon-hTRF1),^3^ and size exclusion chromatography (Superdex columns (Cytiva)).^4^ His_6_-SUMO tag cleavage was achieved via incubation with Ubiquitin-like specific protease 1 (Ulp1) for 1 hour at 25 °C.^5^ For DegP, nickel affinity chromatography was performed under denaturing conditions (6 M guanidium hydrochloride) to remove undesirable bound background proteins. DegP was subsequently refolded via a rapid 100-fold dilution into a refolding buffer (25 mM HEPES, 200 mM NaCl, 1 mM EDTA, pH 7.5) prior to incubation with Ulp1. DegP was then diluted 1:1 in an ammonium sulfate buffer (25 mM HEPES, 2 M (NH_4_)_2_SO_4_, 200 mM NaCl, 1 mM EDTA, pH 7.5), for hydrophobic interaction chromatography.

*CDMS*

CDMS experiments were performed at the University of California, Berkeley, using a custom charge detection mass spectrometer that is described in detail elsewhere.^6^ In brief, ions are formed by nanoelectrospray and are transferred into the mass spectrometer through a stainless-steel ion transfer tube heated to 140 °C. Ions pass through an ion funnel, three RF-only quadrupole ion guides, an ion accelerator, and are trapped in an electrostatic cone trap for 500 ms. A short-time Fourier transform (STFT) is calculated from the time-domain data (1 MHz sampling rate) using a 50 ms window stepped forward in 5 ms increments. From the STFT, individual ion energies, *m*/*z*, z, and mass are obtained throughout the entire time that the ions are trapped.^7,8^ The RF-only quadrupole frequencies and voltages were configured to pass ions with a *m*/*z* greater than ~4,000 for Apo DegP experiments (450 V_pp_, RF 1 at 410 kHz, RF 2 and 3 at 250 kHz) and ~7,000 for DegP/client complex experiments (600 V_pp_, RF 1 at 360 kHz, RF 2 and 3 at 225 kHz). Solutions of ~20 µM DegP and ~10 µM client proteins (in 200 mM ammonium acetate, pH 7) were mixed in equal volume to produce solutions with a 2:1 ratio of DegP to client. Mixtures were stored at ambient temperature (~25 °C) for a minimum of one hour prior to analysis. The electrospray emitter was equilibrated to ambient temperature (~25 °C) for the duration of each measurement unless otherwise specified.

*Variable temperature CDMS*

For variable temperature CDMS measurements, solution temperatures inside the nanoelectrospray emitters were varied using a hollow aluminum cylinder wrapped in resistive nichrome heating wire as described previously.^9,10^ The temperature of the aluminum cylinder was monitored using a J-type thermocouple positioned between the heating wire and the cylinder surface. The temperature was read and logged once per second by a thermocouple reader (Sequent Microsystems) attached to a Raspberry Pi single-board computer. Power was supplied using a 12 VDC power supply. A solid-state relay controlled by a PID loop implemented in Python running on the Raspberry Pi was used to control the temperature. Borosilicate glass emitters were pulled using a P-87 Flaming/Brown micropipette puller (Sutter Instruments, Novato, CA, USA) to an inside tip diameter of 1.7 µm and positioned securely within the bore of the aluminum cylinder such that the emitter tip was approximately even with the end of the aluminum heating block. Nanoelectrospray was initiated by applying ~1.2 to ~1.5 kV to a wire inserted into the electrospray emitter that contacts the solution. The voltage was regularly adjusted to maintain a constant current of ~20 – 40 nA. To confirm that the solution temperature was in equilibrium with the measured block temperature, temperature melts of cytochrome *c* were performed using this same heating device coupled with a Waters quadrupole-time-of-flight (Q-TOF) Premier mass spectrometer (Waters Corporation, Milford, MA). A melting temperature of 72.7 °C from a 10 µM solution (20 mM ammonium acetate, pH ~7) was measured, in agreement with the previously reported value of 73.1 °C,^10^ indicating that the solution temperature and the temperature of the aluminum block are consistent. For the CDMS experiments, the sample solution was allowed to equilibrate to each new temperature for one minute prior to measurements to ensure that the solution temperature reached a steady state. The temperature of the nanoelectrospray emitter was held constant at each specified temperature for 10 min during which time charge detection mass spectra were acquired. Temperature log data and recorded ion data were time-correlated after data acquisition to produce the mass spectra for each temperature. hTRF1 and TrpCage-hTRF1 samples were initially measured at a capillary temperature of 35 °C. The samples were then heated to 50 °C in 5 °C steps, followed by cooling to 26 °C for final data acquisition. IL6-hTRF1 and MNeon-hTRF1 were first measured at 30 °C, then were heated to 50 °C as above, and cooled for a final measurement at 26 °C.

*Circular dichroism thermal melts*

Circular dichroism thermal melts were recorded on a JASCO J-815 spectropolarimeter in 25 mM sodium acetate, pH 7.0, using a 1 mm cuvette. For adequate signal-to-noise, data for apo DegP were collected at 5 µM monomer concentration, IL6-hTRF1 and MNeon-hTRF1 at 10 µM, and hTRF1 and TrpCage-hTRF1 at 20 µM. The circular dichroism signal at 220 nm was recorded during heating from 20 to 90 °C, followed by cooling from 90 to 20 °C at 0.5 °C intervals with a temperature scan rate of 1 °C per minute and a 2-second data integration time.

*Analyses of temperature-dependent circular dichroism data*

Circular dichroism profiles collected as a function of temperature were smoothed using a Savitzky-Golay filter and converted to fraction unfolded, $f_{U}(T)$ (Figure S2), according to

$f_{U}(T)=\frac{(y\left( T \right)- y_{F}\left( T \right))}{(y_{U}\left( T \right) - y_{F}\left( T \right))}$	(1)

where $y\left( T \right)$ is the experimental circular dichroism data, and $y_{F}\left( T \right)$ and $y_{U}\left( T \right)$ are the folded and unfolded baselines. These were assumed to be linear functions of temperature

$y_{i}(T)=m_{i}T+b_{i}$	(2)

where $i\in(F, U)$ and $m_{i}$ and $b_{i}$ are the slopes and intercepts respectively. For each of DegP and the clients, the slopes for the folded and unfolded baselines were estimated to be near 0 from the low and high temperature plateaus of their heating scans; we therefore set to $m_{i}$ to 0 for simplicity and $b_{i}$ to the initial and final values of the melting traces. For a given sample, we found that their unfolded baseline regions for the heating and cooling scans were similar, and we therefore set $y_{F}(T)=y_{U}(T)$ in calculation of $f_{U}(T)$.

*Analysis and visualization of CDMS data*

CDMS data were analyzed using in house Python scripts and Seaborn 0.13.2 kernel density estimate (KDE) features to extract z-average diffusion constants, D_z_. A KDE performs kernel smoothing for probability-density estimation by placing a smooth kernel function on each observation and taking their weighted average, defined for a set of n observations as

$\hat{f_{h}}\left( x \right)=\frac{1}{nh}\sum_{i=1}^{n} K(\frac{x-x_{i}}{h})$	(3)

where $\hat{f_{h}}\left( x \right)$ is the estimated probability density at position *x* (here, the ion mass), *n* is the number of observations (here, detected ions), *x_i_* is the mass of the *i*_th_ ion, *h* is the bandwidth, and *K* is the kernel function. Seaborn employs a Gaussian kernel, given as

$K\left( u \right)= \frac{1}{\sqrt{2\pi}}e^{-\frac{u^{2}}{2}}$	(4)

so that substituting *u = (x-x_i_)/h*, the density estimate becomes

$\hat{f_{h}}\left( x \right)=\frac{1}{nh\sqrt{2\pi}}\sum_{i=1}^{n} e^{\left( -\frac{\left( x-x_{i} \right)^{2}}{2h^{2}} \right)}$	(5)

The bandwidth *h* is a smoothing parameter, related to the standard deviation of the Gaussian kernel centered on each ion through the relation

$h=ah_{0}$; $h_{0}=\sigma n^{\frac{-1}{(d+4)}}$	(6)

where *a* is the bandwidth-adjustment factor, *h_0_* is the automatically selected bandwidth, *σ* is the estimated standard deviation of the ion distribution for a given species, and *d* is the number of dimensions (1 for mass distribution plots, 2 for *m/z* versus charge plots). *σ* and h were automatically adjusted by Seaborn. A smaller value of *a* resolves finer structure, while larger values apply more smoothing. A bandwidth adjustment of 0.1 was applied throughout. The continuous estimate, $\hat{f_{h}}\left( x \right)$, was evaluated on an evenly spaced grid of 1000 points for plotting and fitting. This grid size sets the sampling density of the curve and does not affect the estimate. Two-dimensional KDEs of *m/z* versus charge plots were computed with the same bandwidth and grid setting, rendered as filled density contours, with 50 contour levels and a density threshold of 0.01.

KDE was used here rather than a conventional histogram to avoid the dependence of a histogram on the choice of bin width and bin-edge placement, under which the same data can yield different apparent distributions and features can appear or vanish as artefacts of where the bin edges fall. Replacing the hard binning with a smooth kernel gives a continuous density estimate independent of binning that varies smoothly with the single bandwidth parameter, and is therefore reproducible and well-suited to quantitative curve fitting. The KDE does not increase the intrinsic mass resolution of the measurement or separate physically overlapping species; oligomers of similar mass remain overlapped, as is evident in the distributions. Resolution and quantification of these overlapping oligomeric states were instead achieved by fitting the smooth KDE to a sum of Gaussians described in the following section (see Table S1 for population estimates).

*Quantification of particle fractions and estimation of diffusion constants from CDMS data*

To quantify particle fractions, we assumed that the observed CDMS histograms, $S_{obs.}\left( m \right),$ were sums of the signatures for each apo or client-bound DegP oligomer, $S_{i}\left( m \right)$,

$S_{obs.}(m)=\sum_{i=1}^{N} S_{i}(m)$	(7)

where *N* corresponds to the largest oligomeric state. In what follows, we use *i* as a free index referring to a single state, while *j* is reserved for summation over all states. These are ordinal peak labels and do not relate to oligomer stoichiometry. We described the signatures for individual oligomers as Gaussians given by

$S_{i}\left( m \right)={A_{i}e}^{-\frac{{(m-\mu_{i})}^{2}}{2\sigma_{i}^{2}}}$	(8)

where *m* is the distance in mass from the peak center *µ_i_* that corresponds to the measured effective particle mass, and *A_i_* and *σ_i_* are the peak amplitude and standard deviation respectively. *N* was chosen to be the smallest number of Gaussians that captured the features of the mass distribution, assessed by KDE of the per-ion masses, and that yielded centres consistent with masses expected for DegP oligomers, in multiples of trimers (~140 kDa). For the client-bound DegP samples, we neglected the masses of the bound clients as our CDMS data were unable to resolve client stoichiometries, and with the 2:1 DegP:client ratio used, the additional bound client masses in the cages were negligible compared to DegP (~47 kDa monomer), particularly for hTRF1 (~7 kDa) and TrpCage-hTRF1 (~9 kDa). The masses for client-bound oligomers given here are therefore only approximations. Additional Gaussians were rejected when they did not improve the fit beyond the counting noise or led to fitted parameters with physically unreasonable values. The area under each Gaussian, $a_{i}$, was then calculated according to

$a_{i} = A_{i}\cdot\sigma_{i}\sqrt{2\pi}$	(9)

where *A_i_* and *σ_i_* are the peak amplitude and standard deviation defined in Eq (8). We then assumed that the signal per particle is independent of the oligomeric state, so that $a_{i}$ is proportional to the oligomer concentration *[O]_i_* with the same constant of proportionality for all states. Normalising the area for each oligomer by the sum of the areas then gives the particle fraction

$x_{i} =\frac{a_{i}}{\sum_{j=1}^{N} a_{j}}$ . (10)

Percentages of each oligomer reported in the main text and in Table S1 were calculated by multiplying these fractions by 100%.

*D_z_* values were calculated from CDMS data for comparison with those derived from prior DLS measurements^1,2^ through the following procedure, using in-house Python scripts. Under the assumption above we can also write

$x_{i} =\frac{{[O]}_{i}}{\sum_{j=1}^{N} {[O]}_{j}}=\frac{{[O]}_{i}}{N_{p}}$	(11)

where *N_p_* is the total particle concentration. In order to calculate the set of *[O]_i_* for estimating *D_z_*, *N_p_* must first be calculated using the total monomer concentration

$C_{tot}=\sum_{j=1}^{N} 3n_{j}{[O]}_{j}$	(12)

in which the stoichiometric factor *n* is the number of trimers in a given particle, estimated by dividing the corresponding Gaussian peak center mass, *µ*, by the trimer mass. Rearranging Eq. 11 and substituting into Eq. 12 gives

$C_{\mathrm{tot}} = N_{p}\sum_{j=1}^{N} 3n_{j}x_{j}$	(13)

The sum on the right-hand side is the number-average of protomers per particle,

$\overline{n} = \sum_{j=1}^{N} 3n_{j}x_{j}$	(14)

leading to

$N_{p} = \frac{C_{\mathrm{tot}}}{\overline{n}}$.	(15)

Finally, substituting Eq. 15 into Eq. 11 yields

${[O]}_{i} = \frac{x_{i}}{\overline{n}}C_{\mathrm{tot}}$	(16)

Eqs. 10-16 can be consolidated into a single expression using the fitted areas normalized by their protomer-weighted sum

${[O]}_{i} = C_{\mathrm{tot}}\frac{a_{i}}{3\sum_{j=1}^{N} n_{j}a_{j}}$.	(17)

The set of diffusion constants, *D_i_*, for each oligomer in a given CDMS mass histogram were calculated using a scaling law relating diffusion constants to molecular size.^1,2^ The reference diffusion constant corresponding to a DegP trimer, *D_0_*, was calculated using the known trimer hydrodynamic radius, *r*_3_, of 4.9 nm^2^ and the Stokes-Einstein equation

$D_{0}=\frac{k_{B}T}{6\pi\eta r_{3}}$	(18)

where *k_B_* is the Boltzmann constant, *T* is the absolute temperature, and *η* is the solution viscosity. Viscosities were computed in SEDNTERP.^11^ Because SEDNTERP does not parameterize ammonium acetate, which was used for our CDMS measurements, 200 mM sodium acetate was used as an approximation. The viscosity of the ammonium acetate solution was evaluated at the CDMS measurement temperatures of 25 and 30 °C for calculation of *D_z_* values. The diffusion constants for all higher-order species were then enumerated according to

$D_{i}=D_{0}n_{i}^{p}$	(19)

Where *n* is the number of trimers in a given oligomer, as above, and *p* is an empirically determined scaling constant (-1/3 for spherical oligomers, -0.227 for the oblate path A hexamer).^2^ The oligomer concentrations, masses, and diffusion constants were then combined to calculate *D_z_* values through the relation^1,2,12,13^

$D_{Z}= \frac{\sum_{j=1}^{N} m_{j}^{2}{[O]}_{j}D_{j}}{\sum_{j=1}^{N} m_{j}^{2}{[O]}_{j}}$ .	(20)

Errors for these CDMS-derived *D_z_* values were estimated by a Monte Carlo procedure of 300 iterations. In each iteration, we resampled the per-ion masses with replacement to preserve the counting statistics of the real data. We then recomputed the KDEs of the synthetic resampled distributions and fit them to obtain oligomer concentrations as described above. For each synthetic dataset, the KDE bandwidth was varied by ± 20%, a span that brackets the range over which the KDE resolves the same set of oligomeric states, since narrower kernels fragment populations into noise while broader kernels merge neighbouring oligomers. A given peak’s position was shifted by an amount drawn from a normal distribution with zero mean and a standard deviation of 2% of the peak’s mass, to account for mass-calibration and charge-assignment uncertainty. The trimer hydrodynamic radius, temperature, and the two scaling exponents were also perturbed, using values samples from normal distributions with standard deviations of 5%, 0.5 °C, and 8% respectively, and propagated through the Stokes-Einstein relation (Eq. 18) and the scaling law (Eq. 19) to approximate errors in the diffusion constants for the higher-order oligomers. These values were chosen to allow for systematic error in the trimer radius, which is the dominant term in our error estimates, and to match the 0.4 °C difference between our measured cytochrome c melting temperature and the reported value.^2,10^ The errors in our reported CDMS-derived *D_z_* values are reported as two standard deviations from the mean of the 300 synthetic *D_z_* distributions.^14,15^ DLS *D_z_* values and their associated errors were acquired in a prior study and are reproduced here (Figure 2D and Figure 3D) for comparison with the CDMS values.^1,2^

**Table S1.** Populations for initial and heat-cycled apo and client-bound DegP cages obtained from numerical fitting of CDMS kernel density estimate data.

| Analyte | Mass (MDa) | Oligomeric State | Fractional Area | Error |
| --- | --- | --- | --- | --- |
| Apo, 25 °C | 0.27 | 6mer | 7.5% | 0.85% |
|  | 0.417 | 9mer | 38.5% | 1.5% |
|  | 0.566 | 12mer | 42% | 2.1% |
|  | 0.692 | 15mer | 7% | 1.3% |
|  | 0.832 | 18mer | 3% | 1.9% |
|  | 0.976 | 21mer | 1% | 1% |
|  | 1.09 | 24mer | 1% | 0.5% |
| hTRF1, 35 °C | 0.285 | 6mer | 3.3% | 0.4% |
|  | 0.629 | 12mer | 86.3% | 0.6% |
|  | 1.054 | 21mer | 6.4% | 0.5% |
|  | 1.529 | 30mer | 3.9% | 0.5% |
| hTRF1, Heat cycled up to 50 °C and down to 26 °C | 0.297 | 6mer | 48.8% | 1.6% |
|  | 0.608 | 12mer | 51.2% | 1.6% |
| TrpCage-hTRF1, 35 °C | 0.604 | 12mer | 100% | N/A |
| TrpCage-hTRF1, Heat cycled up to 50 °C and down to 26 °C | 0.331 | 6mer | 36% | 1.6% |
|  | 0.616 | 12mer | 62% | 2.3% |
|  | 0.870 | 18mer | 2% | 1.9% |
| IL6-hTRF1, 30 °C | 0.279 | 6mer | 4.6% | 0.6% |
|  | 0.631 | 12mer | 29% | 1.1% |
|  | 1.014 | 21mer | 13.7% | 0.8% |
|  | 1.454 | 30mer | 52.8% | 1.2% |
| IL6-hTRF1, Heat cycled up to 50 °C and down to 26 °C | 0.27 | 6mer | 76.5% | 7.8% |
|  | 0.422 | 9mer | 15.1% | 8.7% |
|  | 0.614 | 12mer | 8.4% | 2.6% |
| MNeon-hTRF1, 30 °C | 2.422 | 51mer | 1.6% | 0.4% |
|  | 2.917 | 60mer | 6.6% | 1.5% |
|  | 3.412 | 72mer | 14.5% | 1.5% |
|  | 4.340 | 90mer | 77.3% | 1.7% |
| MNeon-hTRF1, Heat cycled up to 50 °C and down to 26 °C | 1.527 | 33mer | 4.6% | 0.9% |
|  | 1.955 | 42mer | 6% | 1% |
|  | 2.415 | 51mer | 5.7% | 4.1% |
|  | 2.892 | 60mer | 11.6% | 1.6% |
|  | 3.361 | 72mer | 29.6% | 1.8% |
|  | 4.286 | 90mer | 42.5% | 1.6% |

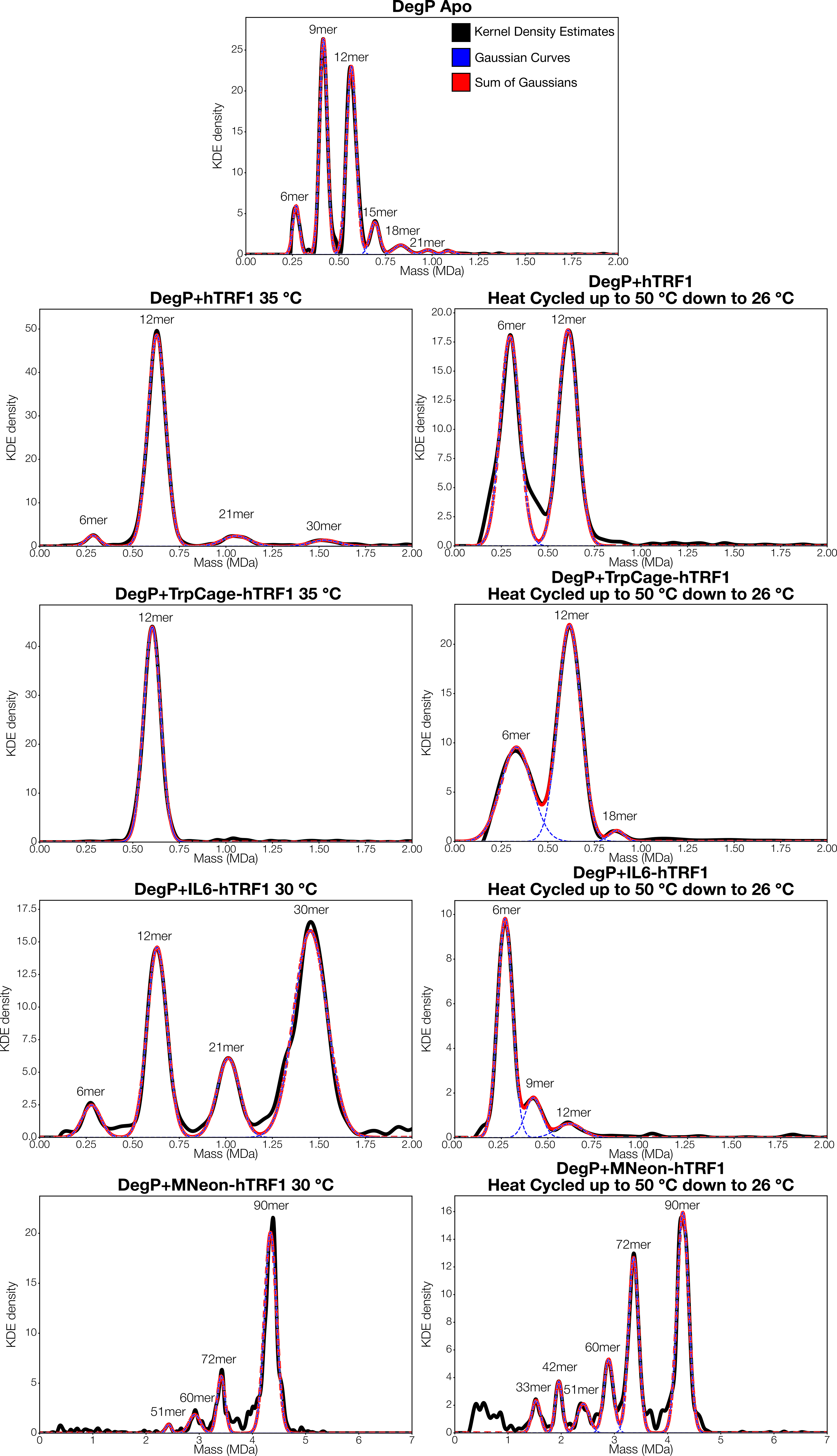

**Figure S1.** Numerical fitting of CDMS kernel density estimates to obtain DegP oligomer concentrations. Kernel density estimates of the experimental data are shown as solid black lines, while fitted Gaussian curves for each oligomeric state are shown as dashed blue lines. The sums of the fitted Gaussians, corresponding to the reconstructed CDMS datasets, are displayed as dashed red lines. Percentages of each state computed from the fitted Gaussians are given in Table S1.

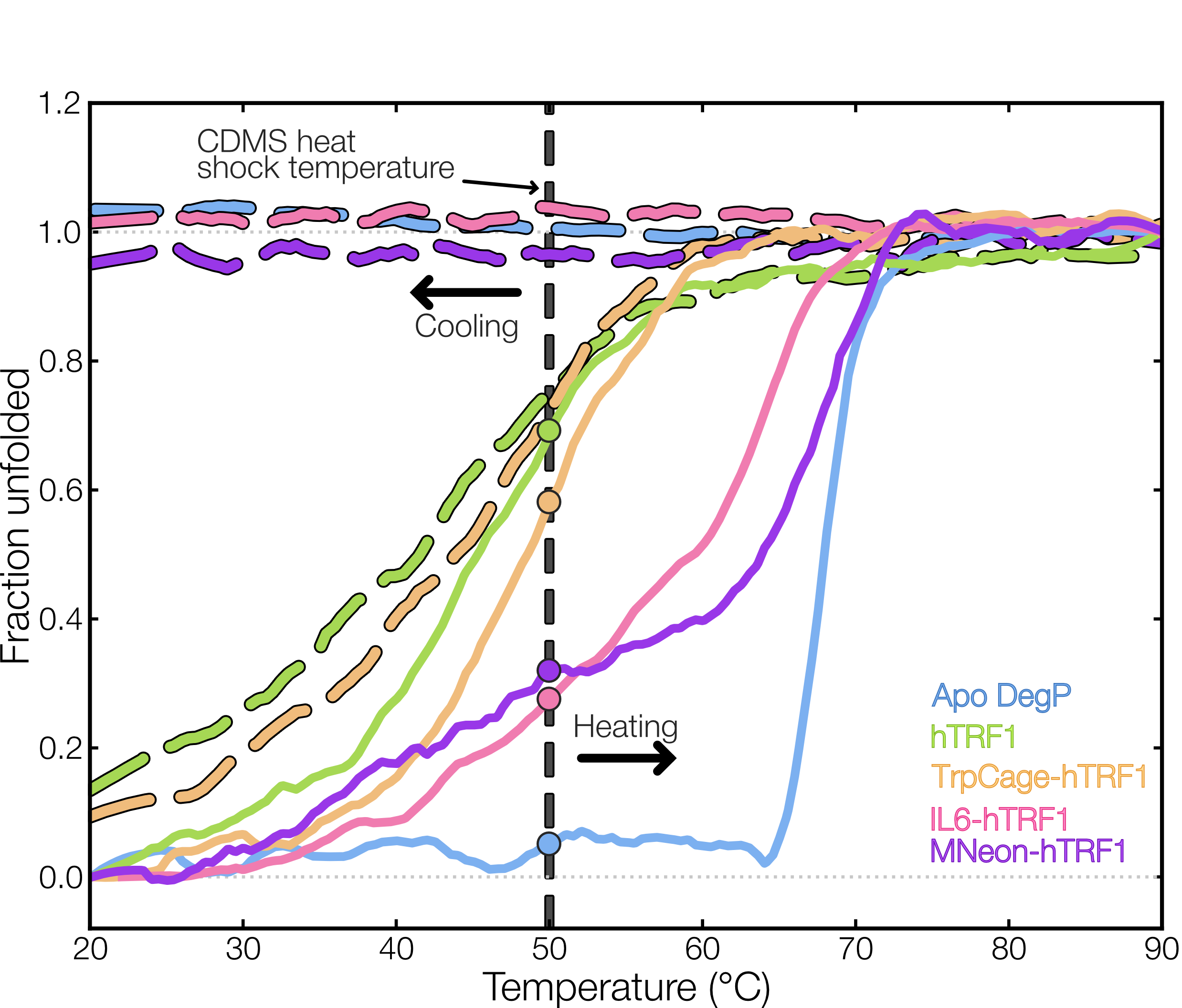

**Figure S2.** Thermal melts of DegP and clients. Fraction unfolded curves obtained from temperature-dependent circular dichroism measurements at 220 nm for apo DegP (blue), hTRF1 (green), TrpCage-hTRF1 (orange), IL6-hTRF1 (pink), and MNeon-hTRF1 (purple) are overlaid. Heating and cooling scans are shown as solid and dashed colored lines, respectively. The dashed vertical line at 50 °C denotes the highest temperature in the heat cycling CDMS experiments. Filled circles mark the fraction unfolded of each client in the heating scan at 50°C. Note that all clients are at least partially unfolded while DegP remains fully folded at this temperature. hTRF1 and TrpCage-hTRF1 partially refold on cooling, whereas apo DegP, IL6-hTF1, and MNeon-hTRF1 have flat cooling scans corresponding to irreversible denaturation.
